# The Rapid Anatomics Tool (RAT): A low-cost root anatomical phenotyping pipeline reveals changes in root anatomy along the root axis

**DOI:** 10.1101/2025.09.05.674482

**Authors:** Dylan H. Jones, Juan C. Baca Cabrera, Dominik Behrend, Darren M. Wells, Joel Swift, Jonathan A. Atkinson, Guillaume Lobet, Meredith T. Hanlon, Hannah M. Schneider

## Abstract

Root anatomical phenotyping has become a demonstrably essential part of investigating root physiology and in acquiring a holistic understanding of plant development. However, accessible high throughput methods for root anatomical analysis are still lacking. Here, we present the Rapid Anatomics Tool (RAT), a novel, low-cost system for high throughput root anatomical imaging with a shallow learning curve for obtaining high quality images suitable for comparative analysis across a number of plant species. Its efficiency comes from combining blockface-like imaging and stain-free imaging using near-ultraviolet (nUV) autofluorescence utilising a combination of low-cost commercial equipment, readily available mechanical components, and custom designed and 3D printed tools. Using this system, we investigated the anatomy of mature tissue along the axis of wheat crown roots, revealing a tendency of reduction in vascular complexity (expressed through a reduction in metaxylem number, area, and mean area per metaxylem file) from the basal to the distal region of the root. This study highlights the importance of thorough sampling strategies for investigating root anatomy in relation to organ function and introduces an accessible, relatively high-throughput method to support such research.

## INTRODUCTION

### Root anatomical phenotyping

Root anatomical phenotyping (typically; quantification of the presence, quantity, size, and distribution of different root cells and tissues) has become a mainstay of the integrated approaches used to investigate root functional traits, as well as in understanding development processes and evolutionary trajectories across a range of species and systems [1]. Phenotypic information of root anatomy can be used to understand radial and axial hydraulic conductance [2–4], nutrient capture [1,5], metabolic cost of root production and maintenance [6,7], the capacity for engagement with microbes and symbionts [8,9], and therefore environmental adaptation [10].

Root anatomy has been studied extensively over the course of history of modern science with interspecies differences in cell and tissue patterning in roots described in literature from the 17th century onwards [11,12]. In the early 20th century, the field of comparative anatomy was advanced significantly by Agnes Arber’s thorough surveys focused on the anatomy of aquatic plants, monocotyledonous plants, and grasses [13]. This was followed by the publication in 1953 of Katherine Esau’s ‘Plant Anatomy’ [14], still regarded as the definitive text in the field, and contemporarily, by the work of renowned botanist, cell biologist, and anatomist, Margaret McCully [15–19].

Several recent root anatomical studies in cereals have explored the associations between plant performance under varied environments and anatomical traits [7,20–22]. Using large germplasm panels not only enables observations of the functional properties of anatomical traits, but when integrated with genetic mapping enables association of anatomical traits with specific genes [1,23]. Root anatomical studies have also been used to understand environmental adaptation during domestication and evolution [24], as well as to investigate the range of functions of specific genes and molecular processes [25]. Significant opportunities remain to explore the precise quantitative impact of anatomical variation on root function under varied environmental conditions and to understand the complex genetic networks that govern these structural traits.

Historically, progress in this area has been concentrated within a few research groups, a limitation imposed by a lack of accessible, high-throughput technology. In recent years, the call for a ‘second green revolution’ focused on the ‘hidden half’ [26,27] has driven an increase in exploration of root physiological traits for crop improvement. Rapid technological developments in recent decades have furnished researchers with a suite of new tools, albeit with a relatively low-throughput, to investigate root physiology, capture phenotypic data on root anatomy, and relate this to functional properties, genetic mechanisms, and plant performance. The study of anatomy provides the essential scaffold for understanding biological function, yet to advance the field, we must move beyond simple qualitative observations and broad correlational relationships. For example, anatomical imaging can be integrated with modelling approaches to quantitatively explore how structural traits influence root function across genotypes or environmental conditions [28].

Being able to image roots at cellular resolution is a critical step in data collection and comparative analysis of root anatomy. Typically, this is done with light microscopy, requiring preparation of thin sections of plant tissue, followed by staining and microscope imaging, or alternatively, the use of highly specialized equipment employing visible light and other forms of radiation or particle detection in order to create a micrograph of plant cell structures. Though these techniques are powerful, versatile, and create enriched datasets alongside 2D anatomical images, brightfield microscopy remains the most common and accessible method of anatomical imaging available.

### Tools and technologies to facilitate anatomical phenotyping of root cross-sections

Despite the increasing availability and speed of digital imaging technologies, sample preparation, particularly the consistent production and handling of thin tissue sections, remains a major bottleneck in anatomical phenotyping workflows. Preparation of thin sections of tissue is essential for transmitted light microscopy, and desirable for confocal laser scanning microscopy, affecting light penetration, reflectance of out of plane light, dye penetration, background fluorescence, and sample topology. This requires at least two flat, consistent, and parallel cuts to be made in the sample, then for this delicate slice of tissue to be transferred to a staining solution and on to a slide without distortion.

A range of tools and techniques are available to prepare root cross-sections for anatomical analysis, each with specific trade-offs in costs (hardware and operating), throughput, versatility, and resolution. Traditional hand sectioning using razor blades or scalpels remains a common method due to its simplicity and minimal equipment requirements [29]. However, this approach demands considerable manual skill to produce consistent, thin sections, and section quality can vary between operators and tissue types. To address the need for more reproducible and accessible sectioning, the *Rapidtome* was developed as a low-cost, open-source alternative to conventional microtomes [29]. This device enables rapid and repeatable hand-sectioning of plant tissues, including roots, with minimal training and no requirement for fragile or high-cost components. More precise sectioning can be achieved using vibratomes and rotary microtomes. Vibratomes employ a vibrating blade to cut sections from fresh or fixed tissue [30]. In contrast, rotary microtomes are typically used with samples embedded in paraffin or resin, and produce ultra-thin sections ideal for detailed histological staining and high-resolution microscopy. While these instruments yield highly consistent results, they require specialized training and are generally more time- and resource-intensive, though can be utilised in high throughput workflows [30].

The recently developed Laser Ablation Tomography (LAT) system operates differently as it combines the ‘cutting’ and ‘imaging’ steps in one interface [31]. LAT uses a powerful laser beam to create one cut surface (that can then be serially ablated in blockface fashion) through a sample, this laser near-simultaneously provides illumination or autofluorescence excitation of the ablated sample surface, while images are captured with a camera positioned to focus on the sample in the cutting plane. Typically, LAT relies on an ultraviolet (UV) laser beam, as many components of the plant root cell wall autofluoresce under UV light. However, infrared (IR) lasers have also been utilized [32]. This unique approach of using the laser for blockface ablation, as well as illumination and excitation, negates the need for sectioning and histological staining. These properties make use of LAT extremely fast to acquire high resolution cross section images. The use of ablation rather than sectioning also means large, tough, or brittle samples that may be intractable in systems requiring sectioning, can be cut and imaged with relative ease. Additionally, the use of high precision sample positioning stages enables LAT systems to be used for both two- and three-dimensional data acquisition in both longitudinal and latitudinal orientations. By substantially increasing imaging throughput, this technology makes it feasible to conduct large-scale genetic mapping studies, such as genome-wide association studies, which are critical for dissecting the genetic architecture of anatomical form and function.

Currently LAT is at the forefront of available anatomical phenotyping methods in terms of throughput, suitability in range of sample types, and resolution achievable in both two and three dimensions. Ultimately however, these systems are extremely scarce, expensive to set up, and time consuming to configure safely. Although these characteristics may limit the wider adoption of LAT for root phenotyping, there are several transferable concepts that can be integrated into more traditional anatomical phenotyping pipelines that can provide high throughput for 2D transverse section imaging. Recent advances in 3D printing have enabled the rapid prototyping of custom components for plant phenotyping, including tools that streamline sample slicing and preparation. The plant research community has embraced this technology, leading to the development of numerous guides, devices, and open-source pipelines aimed at increasing throughput and consistency in anatomical imaging. Through careful optimization of each step, it is now possible to significantly accelerate the workflow and enable high-throughput imaging of plant anatomical features.

### Functional significance of root anatomy: The need for accessible high-throughput phenotyping

Combining concepts from the LAT system with 3D printing and off-the-shelf, low-cost digital microscopes, we have developed a system to obtain root anatomical images with high throughput and with sufficient resolution for trait capture. To demonstrate the utility of our system we present root anatomical data from a survey of historic German winter wheat cultivars, investigating inter- and intra-varietal differences in anatomy. Recent studies have shown that modern breeding practices, shaped by high-input and high-density agricultural systems, may have inadvertently selected for genotypes with smaller root systems and more conservative water use strategies [33].

Basal wheat axial root anatomy is typical of the temperate Poaceae [34]. Seminal roots typically emerge with a single or small number of metaxylem located centrally within the stele, and approximately four to seven layers of cortical cell files. Adventitious (crown) roots may show more variability; early emerging roots produced by the lowest stem node(s) may appear anatomically similar to seminal roots, and later produced roots from successively higher nodes may be relatively enlarged, with higher numbers of metaxylem arranged in a polyarch ring more towards the circumference of the stele and higher numbers of cortical cell file numbers [35–37].

Several root anatomical traits have been shown to be under genetic control, including metaxylem area, stele size, and cortex size [23,25,38–40]. Many of the functional effects of different root anatomical traits in wheat have also been characterised in cereals. For example, enhanced drought tolerance is correlated with narrower vessels due to conservation of soil water through reduced transpiration [41,42]. Root cortical aerenchyma, which are air-filled lacunae in the cortex formed by programmed cell death, reduce the metabolic burden of the root tissue [43,44]. Because a root high in aerenchyma requires less carbon, N, and P for maintenance, these resources can be remobilized for greater root growth, which in turn facilitates more effective soil exploration and resource acquisition [45,46]. Another anatomical trait, fewer files of larger cortical cells, can also reduce metabolic costs and is associated with improved yields in water-limiting environments [47].

An increasing body of work has highlighted the utility of adaptive root anatomical plasticity (a change in development in response to environmental stimuli) in overcoming edaphic stresses resulting from environmental instability, and so has been proposed as a breeding target for crop improvement [20,48,49]. The plasticity described in these studies largely considers the variation in developmentally comparable roots on different plants that have experienced different environments. A further plastic characteristic is the extent of change (and rate of change per unit length) within a single root during its development [50]. Significant anatomical changes along the wheat root axis have been observed, and the amount of change shown to be affected by environmental conditions [50]. We consider these anatomical changes along the root axis to be a critical and largely unexplored dimension in understanding the functional role of root anatomy in the environment. Investigating plastic responses and evaluating their functional effects requires the ability to rapidly image large numbers of roots, quickly, consistently, and with sufficient image quality to reliably quantify anatomical traits. This need only grows when considering the opportunities presented in phenotyping root anatomy in greater resolution along the root axis.

We present an anatomical imaging pipeline that uses commercially available equipment in combination with custom designed 3D printed components that facilitates high throughput anatomical imaging of root tissue. We consider this Rapid Anatomics Tool (RAT) an accessible alternative to more costly anatomical imaging equipment configurations. It provides a pared back toolset well optimised for cross sectional imaging, capable of capturing root anatomical images of comparable quality to leading quality systems that can be used for quantitative analysis. This system was validated by investigating anatomical variation in diverse germplasm. We believe our Rapid Anatomics Tool pipeline to be innovative and functionally distinct from existing methods due to: 1) the integration of the sectioning apparatus with the imaging stage, 2) stain free imaging, and 3) the low cost of and high throughput of the system.

## MATERIALS AND METHODS

### Overview

The Rapid Anatomics Tool (RAT) is a method designed for fast, high-quality anatomical imaging. Its efficiency comes from combining two key principles: blockface-like imaging and stain-free imaging using near-ultraviolet (nUV) autofluorescence.

### Blockface-like Imaging

Our method is inspired by blockface imaging, a technique where a sample, typically held in a supporting matrix, is cut to remove a layer and expose a new, internal surface [51]. The image is then captured directly from this freshly exposed face of the remaining sample block. Typically, blockface imaging is used in conjunction with serial sectioning to obtain 3D structural information. This approach is also used in techniques like Laser Ablation Tomography (LAT) to generate 3D structural information. This differs from many cross-sections anatomical imaging methods where the part of the sample imaged is the thin section cut away from the bulk of the sample, which is often done so that the section can be stained or for transmitted light microscopy. With the RAT system, the sample is positioned in a holder, precisely cut to reveal a crisp and clear cellular face, and this exposed surface is imaged.

### nUV autofluorescence imaging

Several high throughput root anatomical imaging pipelines (including LAT) make use of natural autofluorescence of root cell walls under ultraviolet (UV) or near-ultraviolet illumination (nUV) [52–56]. In many sample types, this can replace the need for staining with chromatophore or fluorophore dyes, reducing the steps required between sample collection and imaging.

By integrating a rapid blockface imaging workflow with stain-free nUV autofluorescence, the RAT system achieves a unique combination of high speed, high quality, and high throughput for anatomical studies. While the core principles of direct blockface imaging and nUV autofluorescence are versatile, in this paper, we present a specific suggested setup. These concepts can be adapted for a variety of equipment and imaging systems to study root anatomy and could potentially be extended to the rapid anatomical analysis of other plant organs.

### Experimental and Technical Design

We present a guide for assembly of the equipment for (Figure 1) and utilization of an anatomical imaging pipeline. This uses custom designed tools and accessible commercially available equipment to enable a hand slicing based workflow to achieve rapid and quality root anatomy image acquisition. Full details of hardware specifics, supplier, and prices are given in the supplementary materials and methods.

**Figure 1:**
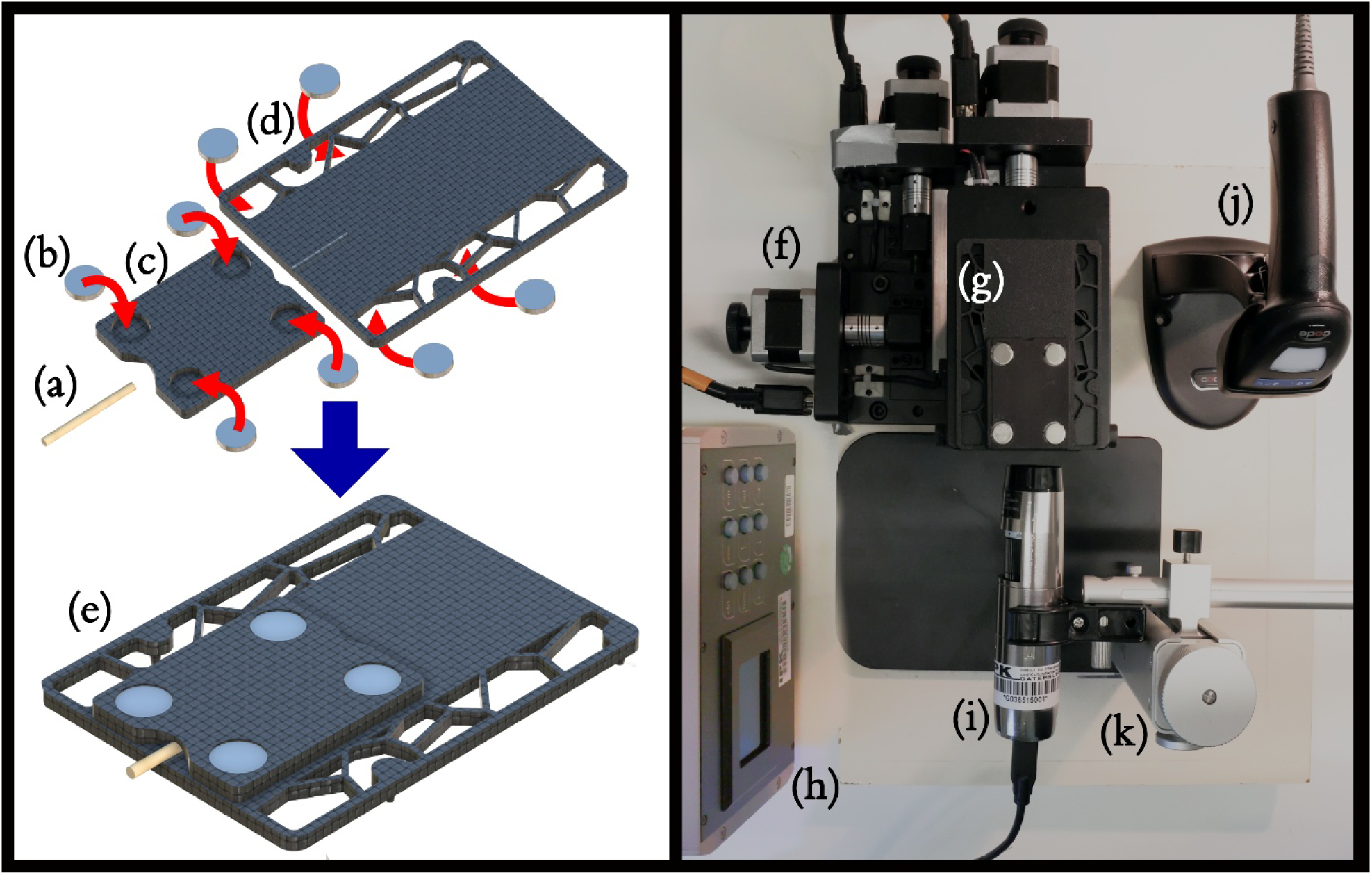
Rapid Anatomics Tool Hardware Assembly Left 3D printed sample holder components (black) magnets (grey) and root piece (beige). Magnets to be held place with superglue in the inset voids. Right, Top-down image showing full hardware setup used in this manuscript. (a) 2 cm root segment, (b) 2 mm x 10 mm disc magnet, (c) 3D printed lid plate, (d) 3D printed base plate, (e) complete RAT root holder assembly, (f) motorized positioning stage, (g) complete RAT root holder assembly, (h) motorized stage controller, (i) USB microscope, (j) 2D barcode reader, (k) USB microscope holder

### Imaging hardware

In our protocol, we use an 8-megapixel USB microscope, with variable magnification 1 X to 220X, with an inbuilt nUV (375 nm) ring light surrounding the objective (Dino-lite Edge AM8517MT-FUW), connected to a standard windows desktop PC. To hold the microscope in place, we use the holder from the microscope manufacturer, though a lab retort stand would function similarly. The manufacturer also provides a software development kit (SDK) that can be used to create custom software control solutions.

### Sample positioning stage

Fine control of sample position is an important element in imaging microscopic structures, in centring the subject of the image, as well as bringing the subject into the correct focal plane. Motorized and computer-controlled stages enable reproducible and precise small movements, and create the opportunity to automate aspects of the sample location process. We have optimized our system around the use of an XYZ motorised control stage, comprising two stacked linear actuator stages with a 50 mm travel range, with a motorized lab on top, these three axes are connected and controlled by a motion controller. The stage manufacturer also provides a software development kit, enabling custom workflows and control of the stages over serial port as well as through their software. Alternatively, we have found the manual positioning stage available from the microscope manufacturer works well and provides sufficiently easy fine manipulation of sample position for use across the full range of magnification levels, and Z axis adjustment can be provided by the microscope stand. Further, many multi-axis fine positioning stages are available commercially or can be manufactured using 3D printing. In order to be amenable with our suggested workflow, a stage should have mounting holes on the top surface, be made of a non-magnetic material, for the mounting holes to align perpendicular to the base, that the base aligns well with the microscope holder, and that the surface of the multi-axis positioning stage have an area greater than 4 cm by 6 cm.

### Sample holder concept and production

The throughput of the LAT system is in part achieved due to the consistency in position of interchangeable samples and the cut surface thereof to be imaged relative to the imaging equipment. To achieve this in our system, we designed a sample holder and manual sectioning jig, intended to achieve the following objectives: 1) to hold a sample of root tissue of a range of sizes sufficiently securely for it to be cut in place without sample movement within the holder, without causing mechanical damage of the region to be imaged; 2) to facilitate cutting of the sample to a consistent fixed protrusion from the sample holder; 3) to locate consistently in a set position relative to the imaging equipment without affecting quality of the image of the sample.

The sample holder was designed for 3D printing in two parts, a base plate with conical ‘feet’ that locates to the mounting holes on the sample stage, and an attachable lid, between which the root sample to be cut and imaged is sandwiched (Figure 1). The sample holder base was designed with a grooved indent in the top surface to help align the sample with the midpoint of the plate consistently. A variety of lids were designed with corresponding recesses with a range of depths to accommodate different diameter roots. The base plate was designed to sit on top and cover the sample positioning stage, and for the sample (before cutting) to overhang the front edge of the base where it would face the camera, we found a size of 7 cm by 12 cm convenient to this end and easy to handle. Lids were designed so the front edge of the lid sat 1 mm recessed from the front edge of the base when the magnets were aligned. Lids were designed to clamp a rear portion of root tightly in place, and for a forward portion of the root to be unclamped and undistorted. As 2 mL microcentrifuge tubes are (in the experience of the authors) the most common vessel for storage of samples for root anatomical imaging, and these typically have an internal depth of approx. 3 cm, lids were designed to hold roots of that length, with 5-10 mm protruding from the front of the base and lid plate sandwich.

On opposite sides of the lid and the base, 2.4 mm deep by 10.4 mm wide cylindrical voids were left, to accommodate 10 mm by 2 mm disc magnets. (A note on assembly: magnets were fixed in place using cyanoacrylate superglue, and we advise preparing the base first, one magnet at a time, then each lid separately, using the base to hold the magnets in the lid plate in place.)

The printer used to produce the sample holders was a Bambu Lab X1C. All settings used were default recommendations of the printer management software per plastic type used. Best results were achieved by printing the base plate in black PLA-CF filament, a polylactic acid-based filament with 5-10% by weight carbon fibre composition. This is abrasion resistant, prints with minimal layer lines, non-fluorescent under UV lighting, matte and non-reflective, suitable for sanding, adheres well with superglue, and prints with a low degree of shrinkage meaning modelled components came out true to size. The lids also printed well in PLA-CF for the same reasons, though for large samples (>6 mm diameter) additional styles of lids were designed to avoid compressing any part of the sample. This was not usually an issue with the small to moderate (0.1-6 mm) diameter roots, but with larger samples, compression of one region occasionally resulted in guttation causing fluid build-up on the cut surface. In these cases, using holders printed using TPU (thermoplastic polyurethane), a soft and more flexible plastic, worked well. The optional blade holder (supplementary materials and methods), can be printed using any rigid plastic.

This is the equipment configuration as used and recommended by the authors, however exact shape and material of the sample holders, assembly of sample positioning stage, and even model of microscope can be amended as to requirements or availability.

### Biological sample preparation

We have used this method or minor variations thereof to successfully section and image thousands of root cross sections for analysis; with sample sizes ranging from 100 µm to 20 mm in diameter. These samples have come from plants grown under varied growth conditions ranging from roots excised from freshly germinated seeds to in-vitro tissue culture, hydroponic culture, compost and sand in controlled environments, as well as from field conditions. Thus far we have found that mature root tissue, outside the root tip and elongation zone, yields the best results, and that image quality diminishes close to the apex. Furthermore, while samples could be processed as fresh tissue, we have noted a tendency towards an increase in image quality after samples were stored in preservative solution for a period of time. For this typically we have subsampled a length of root 2-3 cm long, and stored this in a 2 mL microcentrifuge tube. Tubes are labelled with the corresponding sample name printed on an ethanol resistant sticker label, featuring a 2D Quick Response (QR) code graphic encoding the sample name on the label.

To preserve the sample, we have used solutions of ethanol, ranging from 45% to 70% v/v (depending on availability, transportation needs, and duration of storage intended) filling the remainder of the volume of the tube, and found that a period of one to two weeks at 4 °C yields samples with higher lumen to cell wall contrast than fresh tissues. Roots could be stored for longer periods of time than this (we have not been able to observe age related deterioration in anatomy even in samples stored over 6 months), but 2 weeks was typically sufficient depending on sample size.

Starting with a clean (i.e. free of soil and growth medium) 3 cm length of root, having been stored in ethanol solution for prerequisite time, sample preparation is then straightforward. The root sample is removed from the ethanol solution, handled by the extremities of the sample, and gently blotted on tissue paper (dust free technical wipe paper so as to not transfer UV fluorescent particulate to the sample). The sample is then positioned on the sample holder base plate on top of the indented groove at the front end of the plate with 5 to 10 mm overhanging from the front, making sure to orient the overhanging region perpendicular to the front of the sample holder (Figure 1). An appropriate lid plate can then be selected with a sufficient recess to hold the root firmly without crushing the region at the front of the holder. While holding the base plate *in situ*, the lid plate can then be placed on top of the root.

### Sample Cutting

With the root clamped firmly in place, the root can be cut to length and a fresh surface exposed for imaging. The objective is to obtain a clean cut with minimal damage to the sample and for high throughput processing, for the cut surface to be in a consistent position relative to the sample holder, so minimal sample repositioning and focus adjustment is required.

To achieve this, roots are cut in line with the front of the sample holder base plate. For small to medium sized roots (0.1 to 4 mm), roots can be easily cut by running a Teflon coated single edge safety razor over the front of the holder assembly. Holding the edge of the razor flush with the base plate, and at a 45-degree angle bringing this across the sample with a slight downward motion we found normally gives a clean cut, though this can be adjusted to individual preference. For thicker roots, a thinner blade i.e. half a double-sided razor blade can achieve better cuts. As these can be hard to hold and use loose, we designed a 3D printable razor holder (supplementary materials and methods). The frequency with which the blade will need to be replaced depends on the toughness and size of the samples being processed, for thick field grown roots this may be as frequent as every ∼5-10 samples, and thinner and or artificial media grown roots every ∼20-40 samples. Once the sample is in the holder and cut to length, the holder unit can be mounted in front of the microscope on the sample positioning stage.

### Staining

By using nUV illumination, the need for further contrast staining is eliminated for a large range of root sample types, particularly mature Poaceae axial root samples. This method however is less suitable for visualising tissues that have less secondary thickening of their cell walls, and typically those of a younger developmental age.

To expand the potential range of uses of this setup, we tested several other sample types, and found that the autofluorescence in the cortices and the contrast between the cell walls and the lumen in these tissues were poor. To overcome this, we found a short, direct application of 0.5 µL aqueous solution of fluorescent brightener 28, 0.3 mg mL^-1^ to the cut face, letting this sit for approximately 10 s (adjusted to specific samples where needed), followed by gentle rinsing of the sample in the holder with water and light blotting of the sample holder with tissue paper to remove excess water, enough to improve the fluorescence of the sample to enable imaging. This required readjustment of the image capture settings to account for the change in type of fluorescence, and results in a partial loss of the differential fluorescence between secondary thickened and non-secondary thickened cells, but did enable previously poorly visible cells to become clear.

### Image capture

The digital microscope camera is provided with software (Dino-Capture) of which two versions exist (as of date of submission) (2.0 v 1.553B, and 3.0 v1.1.1.3). We have found version 2.0 more robust and customisable in our use cases, with the ability to add keyboard shortcuts useful to achieving high throughput imaging. That said, we felt that with our specific use case of trying to capture images in high throughput for analysis, improvements could be made. There is a software development kit (SDK) available for the microscope, available for use with LabView (National Instruments) application development software, as well as suitable for use with Python programming language, among others.

To reduce software control steps in our imaging pipeline, we produced an application to be run in Python, displaying a live camera feed, giving the ability to adjust camera parameters and image capture triggered by text entry. Our recommended workflow includes use of a 2D barcode reader to scan the label on the sample tube; with this system image capture is triggered by text entry followed by a 0.1 second pause, saving the image with the scanned code text or entered text as a TIFF file, with the metadata of the image field of view, pixel size, and current magnification level, saved as metadata in the image. The digital microscope camera settings can be adjusted ad hoc during imaging as required by the sample type.

### Plant samples

The RAT system has been tested on a range of diverse plant samples, across multiple devices and set-ups. These tests produced sufficiently clear and high-quality images for anatomical analysis, with high visibility of all cell types. Here we use the system to image and phenotype wheat root anatomy on two winter wheat cultivars, investigating inter- and intra-varietal differences in anatomy.

### Root sampling

Plant growth, management, and root sampling is described in Baca Cabrera et al. 2025 [33]. In brief, a slightly modified version of the wheat root excavation method described by York, 2018 [57] was used. Sampling took place at the end of the tillering stage (BBCH < 30) during two growing seasons, specifically in spring 2023 and spring 2024. All winter wheat cultivars were sampled simultaneously during single-day campaigns. A representative area within each plot was excavated, targeting the top 20–30 cm of soil and covering a similar diameter (around 30 cm). Due to the high planting density, each collected sample typically included 5–10 plants. The excavated plants were placed into sample bags, transported to the laboratory, and kept at 5°C until they could be processed. In the lab, the samples were first soaked in water and then carefully washed. Root crowns were then separated from the shoots near the base, leaving approximately 3 cm of tiller shoot tissue attached, and stored again at 5°C in a solution of water (37.5%), ethanol (37.5%), and glycol (25%). For further analysis, 4 plants per plot were preserved.

### Subsampling for anatomical imaging

Four of these preserved roots per plot were stored in a 30 mL vial in the ethanol, water, glycol solution until subsampling. Each root was dissected into three 2.5 cm lengths, a basal (the 2.5 uppermost cm of the root), middle (from the middle of the root sample, typically 10-12 cm from the base), and a tipwards sample (2 cm towards the middle from the distal most region present, typically 20 cm from the base and or 3 cm back from the root tip). It should be noted that the samples usually did not include the tip, due to the roots having been excavated from the field. These samples were transferred to a 2 mL Eppendorf microcentrifuge tube, and the remainder of the volume was filled with 70% ethanol (v/v) solution. Each root from each pooled plot sample was assigned its own identifier number, and each root position was categorized separately. Sample tubes were labelled with a QR code giving the sample name. Samples were stored at 4 °C in the ethanol solution for a minimum of two weeks.

### Anatomical imaging

Roots were prepared and imaged as described above N=36 per genotype and sampling position. Root samples were largely consistent in diameter, so the digital microscope was used at maximum magnification (220x) for the whole sample set. A small number of samples (< 2 %) had poor cortical autofluorescence, so the staining procedure was used in these cases, this did not affect measurement.

### Analysis

#### Image measurement

Images were measured using FIJI (ImageJ distribution) [58], stele and cortex diameters were measured at their narrowest points using the line tool, areas were measured manually using the polygon tool. For metaxylem, images were converted to 8-bit, and a 2-pixel radius gaussian blur was applied. The magic wand tool was set to a threshold of 20 (adjusted ad-hoc) to select the area of each metaxylem lumen individually.

#### Data processing and analysis

Image data was processed in Microsoft Excel (2019) to create summary data for each image, for each cultivar, and each root position. Yamauchi *et al.* [10] used the ratio between different tissue types within the root to explore diversity in adaptation of Poaceae across an environmental water availability gradient. We have used two of these metrics (xylem to stele ratio, and cortex to stele ratio), multiplied to give a single value (XCS, (Xylem area x Cortex area)-i-Stele area 2) which we found served to aid in identification of trends in change of the proportional relationship between tissues comprising the root cross-sectional area. Data was analysed and statistical comparison of anatomical traits between cultivar and position was carried out with two-way analysis of variance (ANOVA) with Tukey’s multiple comparison correction in Prism 10 (GraphPad) P=0.05.

## RESULTS

### Efficacy of method

The described image acquisition pipeline using the rapid anatomics tool is suitable for imaging a wide range of plant samples at sufficient resolution to enable quantification of cellular anatomical traits. We have used this method to image thousands of root sections for analysis, from species including wheat (*Triticum aestivium*), sorghum (*Sorghum bicolour*), maize (*Zea maize*), cassava (*Manihot esculenta*), and gamagrass (*Tripsacum dactyloides*), and have validated its application with stems, leaves, petioles, and storage roots, from different organ ages. Representative images showing some of the range of species and sample types are shown in Figure 2. This system as described is well optimised for capturing single cross sectional 2D images from a sample in high throughput, and with a low hardware and running cost as well as a shallow learning curve. With just a small amount of practice, researchers without prior experience of anatomical imaging or microscope use can acquire high quality cross section images, and with greater practice, achieve high throughput (∼45 samples per hour for a single user) in doing so.

**Figure 2:**
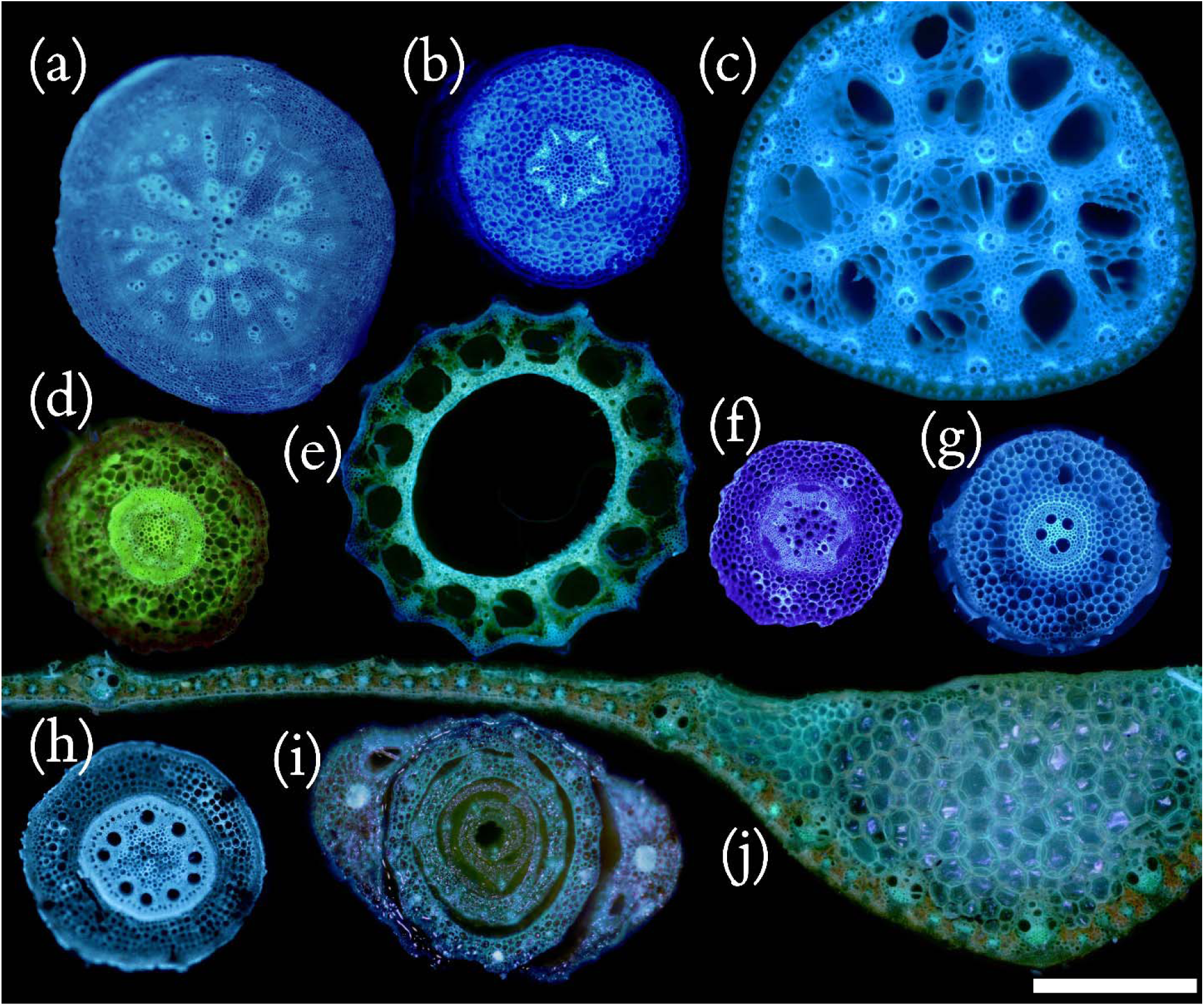
Anatomical images of transverse sections through various plant organs captured using RAT pipeline; (a) horseradish (*Armoracia rusticana*) primary root, (b) Cassava (*Manihot esculenta*) adventitious root, (c) papyrus (*Cyperus papyrus*) stem, (d) Saxifrage (*Saxigraga stolonifera*) stolon, (e) Horsetail (Equisetum) stem, (f) Pea (*Lanthrus oleraceus*) adventitious root, (g) Wheat (*Triticum aestivum*) adventitious root, (h, i, j) Sorghum (*Sorghum bicolor*), (h) adventitious root, (i) seedling stem, and (j) flag leaf; Scale bar 2 mm.

### Root anatomical changes between historic and contemporary wheat cultivars

As a proof of concept, root anatomy was evaluated in crown roots of two German winter wheat cultivars (*T. aestivum L*) one historic (S. Dickkopf, released in 1895), and one contemporary (Tommi, released in 2002), in multiple positions along the root axis. This analysis revealed significant differences between cultivars, as well as significant differences in anatomy along the root axis, and difference in the propensity for change along the root axis between the cultivars.

### Root anatomy differs between cultivars

Significant differences in root anatomical traits were present between cultivars at all three subsampling positions along the root axis. In all instances of significant and non-significant difference, tissue cross sectional area was greater in Dickkopf than in Tommi (Figure 3). In the basal region of the root, this was only significant for metaxylem total area, and in all other sampling regions, this was significant for all cortex, stele, and total metaxylem area.

**Figure 3:**
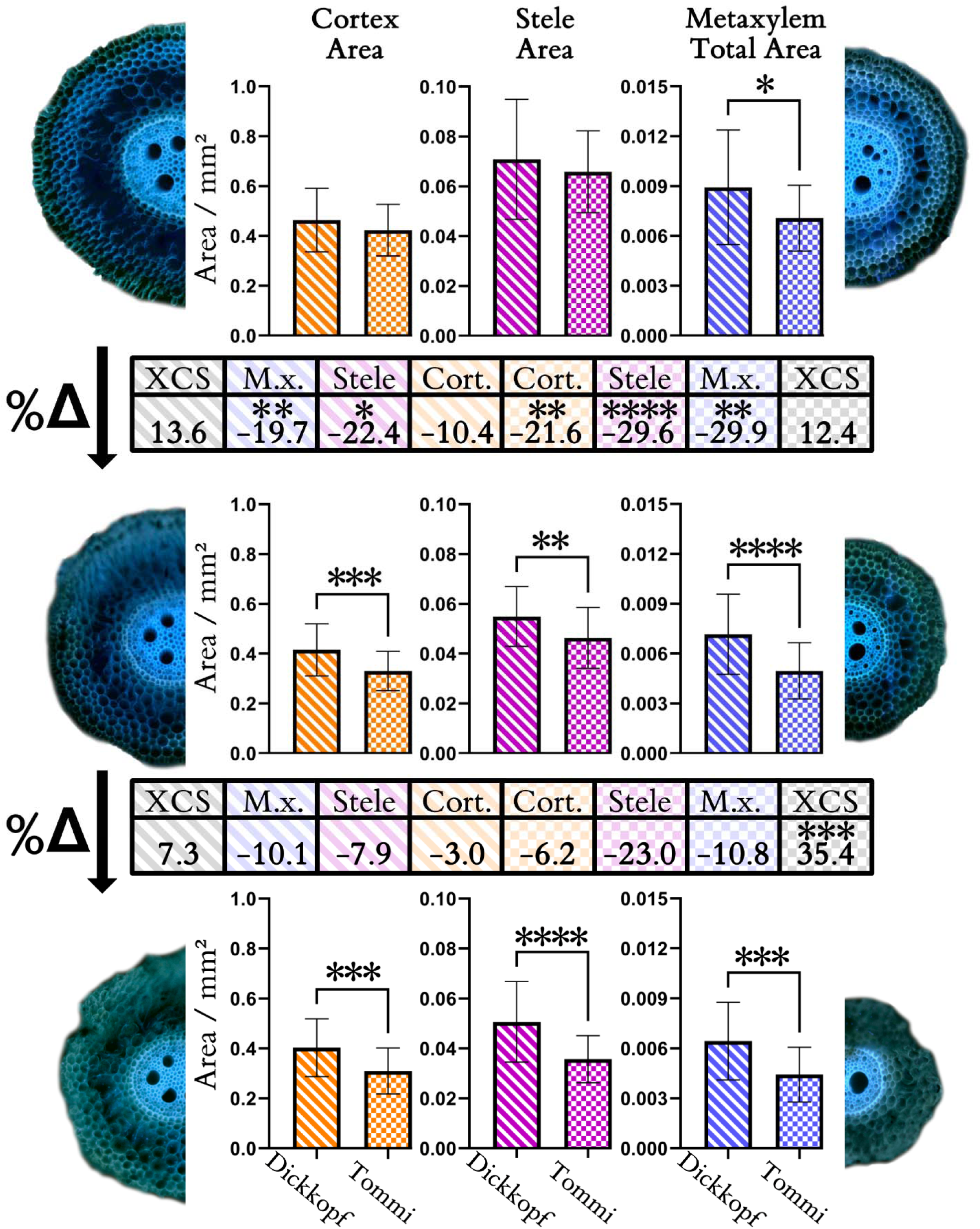
Root anatomy differs between two German winter wheat cultivars as well as along the root axis within cultivars; Root images and graphs: Left side and descending stripes= Dickkopf (1895), Right side and checkered pattern= Tommi (2002) Top row= Root anatomy in basal root region, Middle row= Root anatomy in mid-root region, Bottom row= Root anatomy in distal root region Orange= Cortex area, Magenta= Stele area, Blue= Metaxylem Total Area. Inter-row tables: Percentage change in mean trait value between positions, Left side and Descending stripes= Dickkopf (1895), Right side and Checkered pattern= Tommi (2002), Cort= Cortex area, Stele= Stele area M.x.= Metaxylem total area, XCS= Ratio descriptor of (Meta)Xylem, Cortex, and Stele areas. Statistics: Two-way anova with Tukey multiple comparison test comparing mean values between cultivars and positions, error bars=standard deviation. * =0.05, **=0.01, ***=-.001, ****=0.0001, n=32

### Root anatomy varies along the root axis

There is a clear trend of a decrease in cross sectional area along the root axis in sampling points further from the root base. In Dickkopf the changes in cross sectional area between positions are only significant in the metaxylem and stele, between the basal and middle sample points. In Tommi, root anatomy changes significantly between the basal and middle positions, reflected in a significant decrease in area of stele, cortex, and metaxylem. Though there is a sizeable further percentage decrease in stele area (−23 %) between the middle and tipwards region, this change is non-significant (Figure 3).

### Cultivars show different dynamics of anatomical transition along the root axis

To understand whether these differences in tissue areas between cultivars and along the length of the root represent a shift from isometric to allometric scaling, we used a combination of tissue ratios shown to be effective in relating anatomy to environmental adaptation (the ratios of metaxylem to stele, and cortex to stele) to derive a single value metric. This analysis shows several further distinctions between the cultivars. Firstly, the modern cultivar, Tommi, undergoes a larger change in root anatomy along the root axis than the historic cultivar, Dickkopf. The magnitude of change in tissue area and XCS ratio between positions along the root was greater in Tommi, than Dickkopf in all instances bar XCS between basal and middle sections. The incidence of significance differences in tissue areas between positions is also higher in Tommi than Dickkopf. The difference between the cultivars in change in tissue ratios between sample points is notable and further highlights the differences between the cultivars. Basal to middle ΔXCS is comparable and non-significant for both cultivars, however between the middle and tipward section ΔXCS is much higher in Tommi and statistically significant (Figure 3). This difference describes a significant shift from isometric to allometric scaling in the dynamics of change along the root axis, from the historic to the modern cultivar.

## DISCUSSION

In this study, we present the Rapid Anatomics Tool (RAT), a novel, low-cost, and high-throughput imaging pipeline, and demonstrate its utility by investigating anatomical variation along the axis of adventitious roots in historic and contemporary wheat cultivars. Our work provides two principal contributions: first, a significant methodological advancement that makes rapid root anatomical phenotyping more accessible; and insights into how root anatomy varies within a single root, revealing distinct patterns between cultivars that reflect potential adaptations to different agronomic systems.

The method presented here offers a rapid, low-cost, and high-throughput approach for imaging (primarily) root anatomical traits by harnessing the natural autofluorescence of lignified and suberized tissues under UV illumination. This approach largely eliminates the need for chemical staining or high-end microscopy equipment, addressing major limitations in throughput and accessibility that often constrain anatomical phenotyping in root biology.

When compared to other digital imaging equipment and workflows used for plant anatomical imaging, the RAT system has several advantages (Figure 4) [1,29,59,60]. These are predominantly that it is cheap to set up and operate, and can achieve very high throughputs (Figure 4). Ultimately, it does have an operating cost (other than electricity) in the use of razor blades, where other systems (LAT) may just require power to operate, but this cost is close to negligible and relatively more than offset by the cost of maintenance of the more costly equipment. We feel the resolution is sufficient for anatomical trait quantification of most axial roots and larger lateral roots as well as several other tissues, but for imaging of especially small samples and those lacking secondary thickening, or subcellular detail, other techniques such as electron microscopy (SEM) or confocal laser scanning microscopy (CSLM) may be necessary [15,24,61]. Our optimisation of this method for low cost and throughput in acquisition of 2D anatomical images comes at the cost of versatility. Confocal laser scanning microscopy (CLSM) and light sheet fluorescence microscopy (LSFM) both have the capacity to be used for 2D as well as 3D imaging, as well as quantitative imaging of fluorescence for detection of specific compounds and fluorescent proteins [62]. These, as well as X-ray microCT are non-destructive, and so can be used for 4D anatomical imaging in time series [63]. LAT imaging has potential to be developed further; validation of LAT images with other analysis approaches shows that fluorescence patterns in LAT images can be related to cell wall composition [21]. The use of LAT for sample surface preparation also means impermeable or hard to cut samples that would be challenging to prepare for CLSM can be imaged easily in both two and three dimensions.

**Figure 4:**
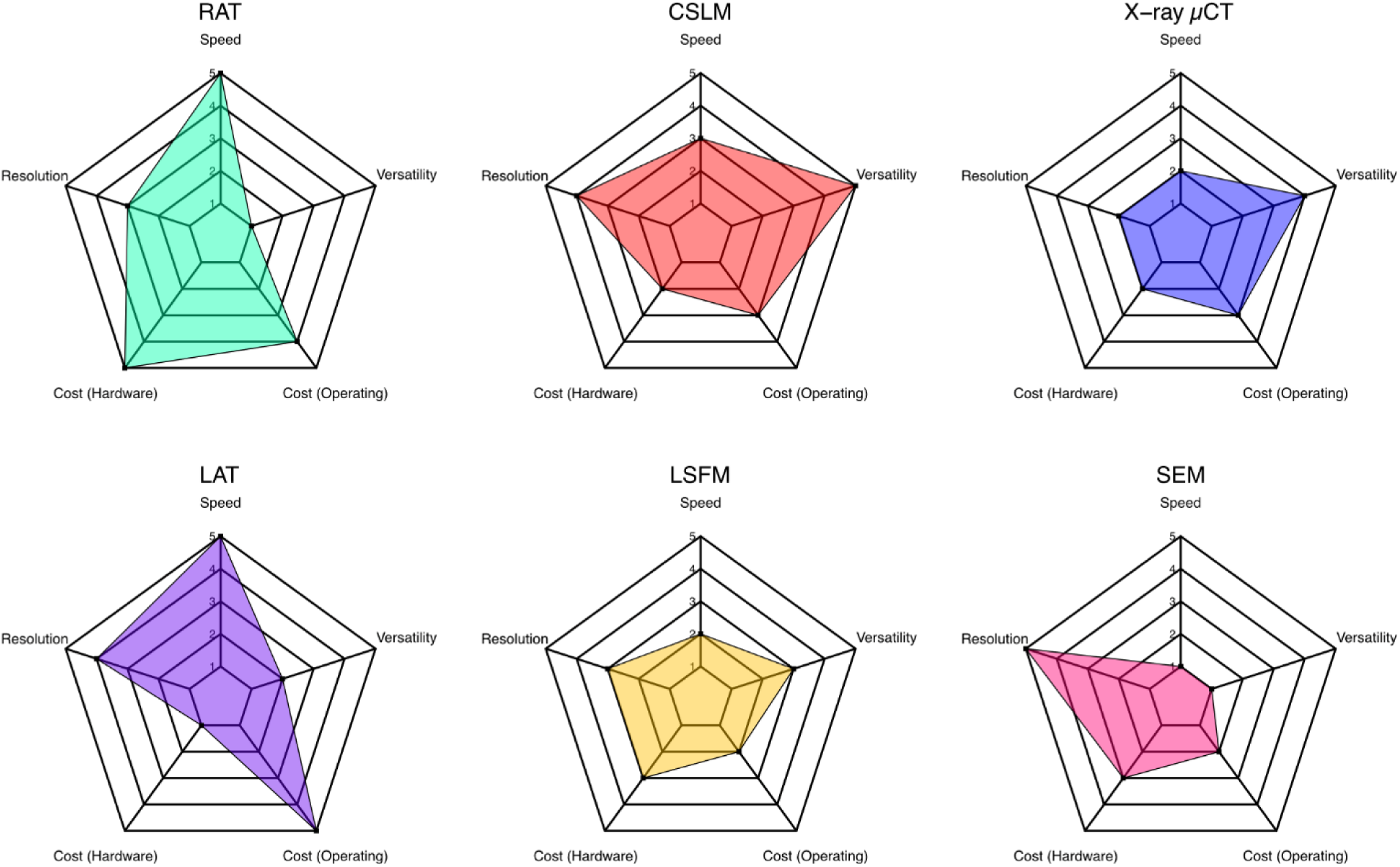
Illustrative plot showing the merits of the described RAT system relative to alternative methods for root anatomical phenotyping. Left to right, top to bottom; RAT: Rapid Anatomics Tool, CLSM: Confocal laser scanning microscopy, X-ray µCT: X-ray micro-computed tomography LAT: Laser Ablation Tomography, LSFM: Light Sheet Fluorescence Microscopy, SEM: Scanning Electron Microscopy. 1, Innermost ring=least desirable (Slow sample preparation, low imaging resolution, expensive hardware, expensive to operate, limited range of applications of the techniques) 5, outermost ring=most desirable (Fast sample preparation and image acquisition, low hardware cost, high micron/pixel, expensive to operate hardware, high application beyond acquisition of 2D anatomical images). Relative merits of key attributes assigned based on reviewed literature [1,15,21,24,29,59,60,61,62,63].

Ultimately though, for our intended purpose of 2D anatomical data collection, our system provides greater or equivalent sample throughput, with little to no sample preparation needed, and at a small fraction of the cost of other systems often employed in anatomical studies (less than four thousand euros compared to tens or hundreds of thousand euros).

### Scope for future improvements

While the RAT system is deliberately optimized for high-throughput phenotyping, it is important to acknowledge its inherent limitations and practical trade-offs. The image resolution is constrained by the consumer-grade optics, which may limit the detection of finer histological details such as the Casparian strip. Furthermore, the reliance on autofluorescence means that image quality can vary depending on the root’s developmental stage, genotype, or environmental conditions. The manual cut through the sample, though effective, may lack the consistency of mechanical or vibratome slicing, which would be preferable for studies requiring the highest precision. Finally, while the mechanical clamp provides essential stability for imaging, it can cause compression damage to the limited portion of the root tissue held within it. Beyond these hardware and methodological constraints, the speed of the workflow introduces important considerations for sample handling. Storing root samples in ∼50% ethanol solution is a double-edged sword; while it prevents microbial decay and clears tissue to improve image contrast, it also makes the samples more prone to drying and cellular collapse upon removal from the preservative. This necessitates rapid imaging to avoid artifacts. Conversely, using lower ethanol concentrations to reduce evaporation can, due to the clamping pressure, result in a guttation effect that floods the cut surface and obscures cellular detail. While alternative preparation methods like critical point drying could potentially solve this, they would significantly increase preparation time and sample brittleness, undermining the system’s primary advantage of speed.

Despite these limitations, the platform’s flexibility in sample type and size enhances its value as an accessible tool for large-scale screening. Our use of computer-controlled motorized stages provides a high degree of precision for tasks like focus stacking, but these are not essential. The core benefits of the system can be achieved with more affordable, manually adjusted stages, including many open-source designs available for 3D printing. Looking forward, there are several clear avenues for improvement. The system could be upgraded with higher-resolution imaging modules, integrated with automated sectioning systems to boost consistency, or coupled with real-time trait extraction using machine learning approaches. In summary, our UV autofluorescence imaging platform, even with its trade-offs, offers an accessible and scalable solution that facilitates the broader adoption of belowground trait analysis in root research and plant breeding.

### Root anatomical differences in variation along root axis

Leveraging RAT to investigate inter- and intra-specific variation in wheat roots, we discovered many root anatomical traits were significantly different between the historic and modern wheat cultivars surveyed. These differences became more frequent and more statistically pronounced when comparing positions on the root further along the root axis from the basal region. The anatomical differences along the root axis in the modern cultivar are most strongly observed in the vascular traits, the stele (−52.6 %) and metaxylem total (−40.7 %) areas, more so than the cortex area (−27.8 %). Similarly, in the historic cultivar, the relative changes in root anatomy along the root axis were more strongly observed in stele (−30.3 %), and metaxylem total (−29.8 %) area than in the cortex (−13.4 %, non-significant) area. Comparing the differences in the ratiometric root composition between sampling positions and cultivar makes these differences especially clear. Using this integrated metric accounting for metaxylem, cortex, and stele area as a proportion of the total root area (XCS) we can see that the root anatomy of the modern cultivar is not just changing along the length of the root, but changing its proportional area by tissue type allometrically. This transition to a different root anatomical tissue composition appears driven by the vasculature in these instances. While there are also differences in the percentage change in area between positions in the historic cultivar, they are less stark than in the modern cultivar, do not represent a significant change in XCS ratio, and therefore can be viewed as more isometric than allometric in nature. These results are interesting to note within the context of the broader root physiological results from the trial that yielded these samples [33]. In this study, it was seen that axial root number significantly decreased from the historic to the modern cultivar, and that measured and simulated whole root system hydraulic conductance also followed this trajectory. Interesting to note too is that in the root system analysis, there was no significant effect of breeding (cultivar release date) on crown root diameter [33]. This is reflected in our anatomical analysis, though the significant differences we observe in the vasculature between the historic and modern cultivar may well contribute to the described difference in root system hydraulic conductance. This highlights the importance of anatomical studies and the accessibility of tools to investigate anatomy in gaining a more complete and integrated understanding of the relationship between root physiology and hydraulic activity.

In the emerging field of high throughput root comparative anatomy, we consider the distinction between isometric changes and allometric transitions along the root axis to be a relatively underexplored phenomenon, potentially in part due to the lack of accessible sufficiently high throughput techniques for root anatomical imaging. While anatomical changes along individual axial roots have been shown to occur in a range of species, such as wheat [36,40,48,64–66], rice [36], millet [67], maize [28,68], wheatgrass, and alfalfa [35]. What drives these changes along the root also bears considering; while not mechanistically investigated, it is suggested that these are likely to relate to changes in Indole-3-acetic acid (IAA) gradient due to increasing distance from young leaves [64]. Additionally, plastic anatomical changes along individual roots have been shown to be affected by environmental factors such as hypoxia and compaction [50]. Whether these hormonal signals and or environmental factors dictate whether or not these changes occur, or the functional implications, rate, and position of these changes remains to be explored.

These findings demonstrate the efficacy of our pipeline; that the image quality is sufficient to observe and quantify variation in key functional traits, and that these images can be captured in sufficient throughput that it is feasible as well as beneficial to expand the scope of conventional sampling strategies for anatomical phenotyping. Relative to many other hardware requirements to achieve comparable image quality and speed (for specifically 2D anatomical phenotyping), our platform is cheap to purchase the components for, and run, representing good value for money, and easy to set up and operate.

## Conclusions

This research yielded two key outcomes: a more accessible method for rapid root anatomical phenotyping using nUV autofluorescence and blockface imaging, and a new understanding of intra-root anatomical variation. This study successfully developed and validated the Rapid Anatomics Tool (RAT), a low-cost, high-throughput pipeline that makes detailed root anatomical analysis accessible to a broader range of researchers. By applying this system, we have provided evidence that significant, allometric anatomical variation exists not only between wheat cultivars but also along the length of a single root. This work underscores the critical importance of strategic, multi-position sampling in root biology and provides the research community with an effective tool to explore the ‘hidden half’ in greater detail, accelerating our efforts to develop more resilient and efficient crops. This development and democratization of advanced imaging and analysis pipelines will be revolutionizing the field. This leap forward will finally enable the broader scientific community to systematically link genes to the anatomical phenotypes that drive physiological outcomes.

## Supporting information

Supplemental File 1: User Guide

Supplemental Materials: 3D Files and Software

## ACKNOWLEDGMENTS

**General**: Thanks to Annegret Wolf and Petra Linow, and GPW group members at the IPK for their feedback on RAT design and usability

## Author contributions

The system was conceived by H.M.S. and M.T.H. and developed by D.H.J. with input from H.M.S, J.S., D.M.W, J.A.A., and M.T.H., plant growth, sampling, and management was carried out by J.B.C., D.B., and G.L., testing and generation of images by J.S., M.T.H., and D.H.J, writing of the manuscript was led by D.H.J., all authors contributed to writing, editing, and approved the manuscript.

## DATA AVAILABILITY

3D design files (STL) are provided in the supplementary material. The Python script used to control image acquisition using the specific USB microscope used in this study is available in the supplementary materials, though requires drivers from the software development kit from the manufacturer. Datasets generated and analysed are available from the corresponding authors on reasonable request.

## SUPPLEMENTARY MATERIALS

Rapid Anatomics Tool Assembly and Use Guide (pdf), RAT 3D Print Files and Microscope Control Script (ZIP file)

