## Supplemental File 1: User Guide for "The Rapid Anatomics Tool (RAT): A low-cost root anatomical phenotyping pipeline reveals changes in root anatomy along the root axis"

### Supplemental Material

#### Rapid Anatomics Tool Assembly and Use Guide

After printing the RAT root holder components and glueing the disk magnets in place in the correct orientation, the holder can be used.

| | | | |
|---|---|---|---|
| Step 1- Components prepared, base plate set out | 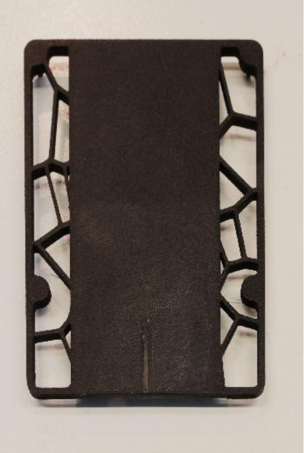 | Step 4- Align razor blade with edge of base plate, keeping the razor perpendicular to the root. This may be best achieved holding the root holder upwards, and holding the razor by its edges | 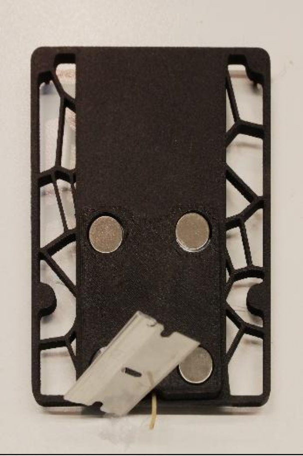 |
| Step 2- Place root on top of base plate. Align sample with central line, approximately 1 cm overhanging the edge of the base plate, with the root perpendicular to edge of base plate | 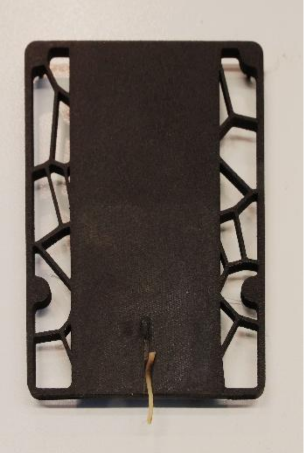 | Step 5- Sweep razor blade through the protruding sample, cutting off excess and leaving cut surface flush with edge of base. A diagonal, forward and downward motion is effective | 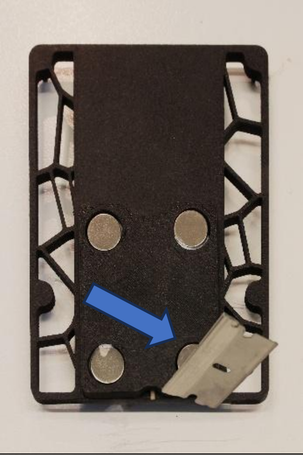 |
| Step 3- Lay appropriately sized lid plate on top of root piece, completing sandwich of root between base and lid. The root should now be secure in the holder | 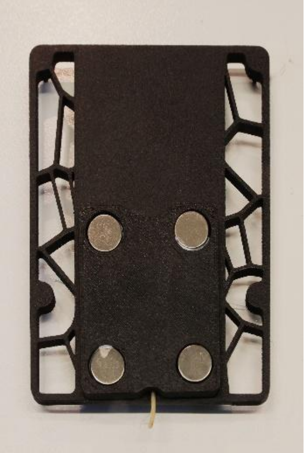 | Step 6- Taking care not to dislodge cut sample, transfer holder to opposite microscope station. If needed, stain can be applied here, or if fluid builds on cut surface, blot gently with dust free tissue paper | 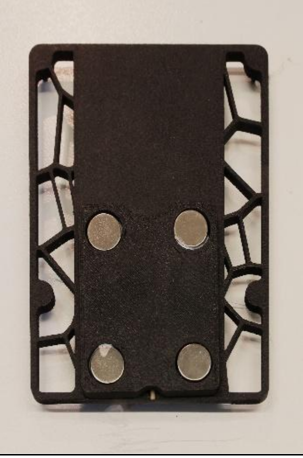 |

#### Razor Blades

The frequency with which blades need replacing depends on the sample properties and care in use. Blade use life can be extended by being careful not to lay the razor on the blade, bring it into contact with hard surfaces other than the sample, and by using different areas of the blade edge instead of replacing the blade. For thin or soft samples, we find the rigid single sided razor as shown in the pictures to be most suitable. For thicker, and tougher samples a thinner cutting edge is useful, such as half of a double edge razor blade. These can be more challenging to hold safely, so we designed a razor blade holder with an elongated handle, recesses for magnets to hold the blade in place, and a bowed region to hold the blade securely between two points (RAT_razor_holder). 2 mm thick 2 mm diameter disk magnets can be glued into the razor holder, which can hold the razor blade in place when this too is glued in place with quick setting superglue (Assembly image below left). The handled blade can then be used more safely and dexterously, in the same way as the single edged razor (below right). When replacement is needed, the razor blades can be easily removed safely with plyers.

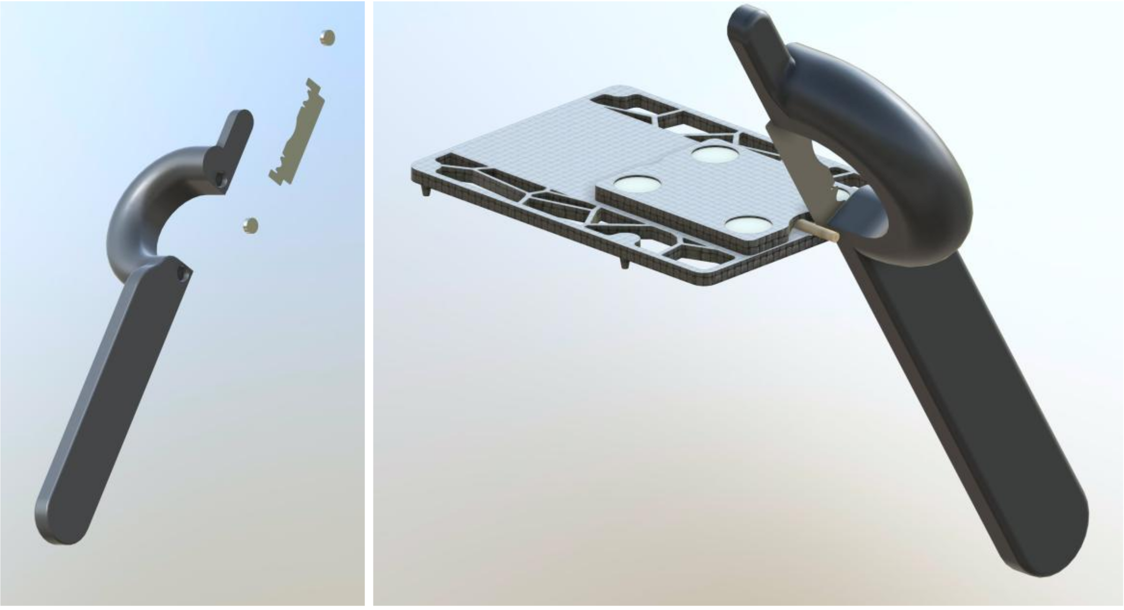

#### Equipment List

| Item | Product | Price | Supplier |
|---|---|---|---|
| USB Microscope | Dino-Lite Edge AM8517MT-FUW | $1,389.00 | Dino-Lite |
| Motorized Linear Stage | Motorized Linear Stage Model: MOX-02-50 | $396.00 | Optic Focus |
| Motorized Lab Jack | Motorized Lab Jack Model: MOZ-80-50 | $294.00 | Optic Focus |
| Motion Controller | Motion Controller (for NEMA17 Stepper Motor, 220V) Model: MOC-01-1-220 | $767.00 | Optic Focus |
| Microscope Stand | Dino-Lite Versatile Precision Mount RK-10A | $299.00 | Dino-Lite |
| Manual Stage | Dunwell for Dino-Lite MS15X-S1 | $169.00 | Dino-Lite |
| Barcode Reader | Brady CR950 Wired Handheld Barcode Reader - 1D, 2D, QR Code | €204.39 | Brady |
| PLA-CF 3D printing filament | BambuLab PLA-CF Black (14100) (1 kg) | €26.99 | BambuLab |

| | | | |
|---|---|---|---|
| Fluorescent Brightener 28 | Fluorescent Brightener 28, powder microbiology stain 4404-43-7, 5g ICNA0215806705 | €272.00 | Avantor |
| Single edge razor blades | Razor Blades, Stainless Steel, GEM, PTFE-Coated, Extra Keen, Washed Version, 115 pieces E71970-WA | €93.65 | ScienceServices |
| Double edge razor blades | Double Edged Razor Blades, Personna, Stainless Steel, PTFE-Coated, 250 pieces E72000 | €116.38 | ScienceServices |

#### Software List

| Name | Function | Version | Supplier |
|---|---|---|---|
| DinoCapture 2.0 | USB Microscope control | 1.5.52 | Dino-Lite |
| Dino-Lite Windows Software Development Kit (SDK) | Custom workflow for USB Microscope control | V3.0.56.6 | Dino-Lite |
| 7SCSCR | Motorised stage controller | 7SCSCR for X64 | Optic Focus |
| Spyder | Python Interactive Development Environment used to control camera with SDK | Spyder 6.0.7 | Spyder-ide |

#### Imaging and Software Control

To use the DinoCapture 2.0 for imaging cut roots as described in the article we recommend a specific setup and range of settings. For image consistency, imaging should be done out of direct sunlight or ambient room lighting, either in a dark room or dark box. Using the Dino-Lite Edge AM8517MT-FUW, lighting setting 2 should be used, at maximum (level 31) light intensity, with a shutter speed of 1/11 seconds, and an iso of 200. Auto-colour balance should be switched to SET, and in the image settings, toggled to off. For our imaging we have used a camera brightness of 170, contrast of 26, and all other settings as default.

#### Custom Software Control

As described, a custom software control pipeline was created in Python using the software development kit requestable from the manufacturer. This requires prior acquisition and installation of the SDK .dll file.
The Python script is named 'RAT_DinoLite_Custom_Workflow_Controller'

This script is functional and optimised for throughput when used with a barcode reader, though offers little in terms of graphical user interface or file management, however can be freely modified to add capacity in these areas if desired given the use case.

Image scales are based on values read from the microscope hardware, and may not necessarily represent exact object size if the microscope has been improperly calibrated prior to use. We recommend minimal adjustment of magnification during imaging of samples intended for comparative analysis, and verification of scales using a fiduciary marker between magnification adjustments.

#### Instructions

When running, this script will open a live view window of the camera feed. The camera setting values can be changed in the scripts 'DEFAULT_SETTINGS' list, which are currently set as optimal for X220 magnification imaging of roots and are as used for data collection in the article. Settings can be changed by selecting the live view window, pressing the 'alt' keyboard button, then using the keyboard shortcuts described below, different settings can be input in the python console, and the settings editing mode can be exited with 'alt'

| Key | Function | Notes |
|---|---|---|
| e | Set Exposure | Prompts for a value (10–20000) |
| a | Toggle Auto-Exposure | 0 = Off, 1 = On |
| f | Show Field of View & AMR | Prints to console |
| i | Set Lighting Mode | 1 = UV, 2 = White, 3 = Both, 4 = Off |
| l | Set Lighting Level | 1–32 |
| b | Set Image Brightness | Integer value |
| r | Get White Balance Range | Prints to console |
| c | Set Contrast | Integer value |
| w | Set White Balance | Integer value |
| h | Set Hue | Integer value |
| s | Set Saturation | Integer value |
| p | Set Sharpness | Integer value |
| d | Restore Default Settings | Uses values in DEFAULT_SETTINGS |
| g | Set Gain | Integer value |
| q | Quit Script | Closes camera and window |

While not in 'alt' mode, the script is in acquisition mode. This is optimised for use with a barcode reader, where a text string is delivered instantaneously. While in this mode the script listens for keyboard events via pynput. Each typed character is appended to a string, and if no new character is typed for 0.5 seconds, the name is considered complete and a TIFF image is captured and saved with the input text as the file name. The script filters characters in the input name, removing invalid characters and replacing them with '_', and appends a timestamp in the format YYYYMMDD_HHMMSS. While saving, the script adds metadata to the file, including pixel resolution (mm), field of view, and the magnification level used on the microscope during capture.
