## Supplemental Materials: 3D Files and Software for "The Rapid Anatomics Tool (RAT): A low-cost root anatomical phenotyping pipeline reveals changes in root anatomy along the root axis": RAT_DinoLite_Custom_Workflow_Controller.py

```
import cv2
import tifffile
import time
import os
from datetime import datetime
from pynput import keyboard
from dnx64 import DNX64

# Constants
WINDOW_WIDTH, WINDOW_HEIGHT = 768, 432
CAMERA_WIDTH, CAMERA_HEIGHT, CAMERA_FPS = 3840, 2160, 60
DNX64_PATH = "Must Be requested https://www.dinolite.us/support/about-the-dino-lite-sdk/?_gl=1*hij2ud*_up*MQ..*_ga*MTgwNzA2OTM3LjE3NTQ3NTQ3ODU.*_ga_5L45SNB5NE*czE3NTQ3NTQ3ODQkbzEkZzAkdDE3NTQ3NTQ3ODQkajYwJGwwJGgw"
DEVICE_INDEX = 0
COMMAND_TIME = 0.1
barcode_complete_timeout = 0.5
SAVE_DIR = "SAVE_DIRECTORY"
os.makedirs(SAVE_DIR, exist_ok=True)

# Initialize microscope
microscope = DNX64(DNX64_PATH)
alt_mode = False
barcode_text = ""
last_input_time = time.time()

# Default preselected values
DEFAULT_SETTINGS = {
    "Exposure": 800, #10-20000
    "AutoExposure": 0,  # 0: Off, 1: On
    "Brightness": 100,
    "Contrast": 30,
    "WhiteBalance": 30,
    "Hue": 0,
    "Saturation": 15,
    "Sharpness": 10,
    "Gain": 2,
    "FrameRate": 30,
    "FLCSwitch": 1,  # 1: UV, 2: White, 3: Both, 4: Off
    "FLCLevel": 20,  # Default intensity level 1-32
}

# Adjustable properties
VIDEO_PROPERTIES = {
    "Brightness": 0,
    "Contrast": 1,
    "Hue": 2,
    "Saturation": 3,
    "Sharpness": 4,
    "WhiteBalance": 7,
    "Gain": 9,
}

# Utility Functions
def apply_defaults():
    print("Restoring default settings...")
    toggle_auto_exposure(DEFAULT_SETTINGS["AutoExposure"])
    set_exposure(DEFAULT_SETTINGS["Exposure"])
    set_video_property("Brightness", DEFAULT_SETTINGS["Brightness"])
    set_video_property("Contrast", DEFAULT_SETTINGS["Contrast"])
    set_video_property("Hue", DEFAULT_SETTINGS["Hue"])
    set_video_property("Saturation", DEFAULT_SETTINGS["Saturation"])
    set_video_property("Sharpness", DEFAULT_SETTINGS["Sharpness"])
    set_video_property("Gain", DEFAULT_SETTINGS["Gain"])
    set_flc_switch(DEFAULT_SETTINGS["FLCSwitch"])
    set_flc_level(DEFAULT_SETTINGS["FLCLevel"])
    print("Defaults applied.")

def set_exposure(value):
    microscope.Init()
    microscope.SetAutoExposure(DEVICE_INDEX, 0)
    microscope.SetExposureValue(DEVICE_INDEX, value)
    time.sleep(COMMAND_TIME)
    print(f"Exposure set to {value}")

def toggle_auto_exposure(state):
    microscope.Init()
    microscope.SetAutoExposure(DEVICE_INDEX, state)
    time.sleep(COMMAND_TIME)
    print(f"Auto-exposure {'enabled' if state else 'disabled'}")

def set_video_property(property_name, value):
    microscope.Init()
    microscope.SetVideoProcAmp(VIDEO_PROPERTIES[property_name], value)
    time.sleep(COMMAND_TIME)
    print(f"{property_name} set to {value}")

def get_video_property(WhiteBalance):
    microscope.Init()
    WB = microscope.GetVideoProcAmpValueRange(VIDEO_PROPERTIES[WhiteBalance])
    time.sleep(COMMAND_TIME)
    print("WB")
    return WB

def set_flc_switch(switch_state):
    microscope.Init()
    microscope.SetFLCSwitch(DEVICE_INDEX, switch_state)
    time.sleep(COMMAND_TIME)
    print(f"FLC Switch set to {switch_state}")

def set_flc_level(level):
    microscope.Init()
    microscope.SetFLCLevel(DEVICE_INDEX, level)
    time.sleep(COMMAND_TIME)
    print(f"FLC Level set to {level}")

def get_fov():
    microscope.Init()
    amr = microscope.GetAMR(DEVICE_INDEX)
    fov = microscope.FOVx(DEVICE_INDEX, amr)
    fov_mm = round(fov / 1000, 2)
    print(f"Field of View (FOV): {fov_mm} mm")
    return fov_mm, amr

def capture_tiff(frame, fov, amr, name):
    rgb_frame = cv2.cvtColor(frame, cv2.COLOR_BGR2RGB)
    pixel_size_mm = fov / CAMERA_WIDTH
    resolution = 1 / pixel_size_mm
    timestamp = datetime.now().strftime("%Y%m%d_%H%M%S")
    filename = os.path.join(SAVE_DIR, f"{name}_{timestamp}.tiff")

    metadata = {
        "XResolution": (resolution, 1),
        "YResolution": (resolution, 1),
        "ResolutionUnit": 3,
        "FOV": f"{fov} mm",
        "AMR": f"{amr}",
    }

    tifffile.imwrite(
        filename, rgb_frame, resolution=(resolution, resolution), metadata=metadata
    )
    print(f"Saved image to {filename} with scale metadata: {metadata}")

def initialize_camera():
    camera = cv2.VideoCapture(DEVICE_INDEX, cv2.CAP_DSHOW)
    camera.set(cv2.CAP_PROP_FPS, CAMERA_FPS)
    camera.set(cv2.CAP_PROP_FRAME_WIDTH, CAMERA_WIDTH)
    camera.set(cv2.CAP_PROP_FRAME_HEIGHT, CAMERA_HEIGHT)
    return camera

def process_frame(frame):
    return cv2.resize(frame, (WINDOW_WIDTH, WINDOW_HEIGHT))

def process_barcode_input(key, frame):
    global barcode_text, last_input_time
    try:
        barcode_text += key.char
        last_input_time = time.time()
    except AttributeError:
        pass

def check_for_barcode_complete(frame):
    global barcode_text, last_input_time
    if barcode_text and (time.time() - last_input_time) > barcode_complete_timeout:
        print(f"Text/Barcode complete: {barcode_text}")
        
        # Sanitize barcode for use in filenames
        safe_barcode = "".join(c if c.isalnum() or c in (' ', '-', '_') else '_' for c in barcode_text)
        if barcode_text != safe_barcode:
            print(f"⚠️  Barcode '{barcode_text}' contained invalid characters. Saved as '{safe_barcode}' instead.")

        fov, amr = get_fov()
        capture_tiff(frame, fov, amr, safe_barcode)
        barcode_text = ""

def start_camera():
    global alt_mode
    camera = initialize_camera()
    if not camera.isOpened():
        print("Error opening the camera device.")
        return

    print("Press 'Alt + [key]' to edit settings. Type to capture image. Press 'q' to quit.")
    listener = keyboard.Listener(on_press=lambda key: process_barcode_input(key, frame))
    listener.start()

    while True:
        ret, frame = camera.read()
        if ret:
            resized_frame = process_frame(frame)
            cv2.imshow("Dino-Lite Camera", resized_frame)

        if not alt_mode:
            check_for_barcode_complete(frame)

        key = cv2.waitKey(25) & 0xFF

        if key == 18:
            alt_mode = not alt_mode
            print(f"Alt mode {'enabled' if alt_mode else 'disabled'}.")

        if alt_mode:
            if key == ord('e'):
                set_exposure(int(input("Enter exposure value: ")))
            elif key == ord('a'):
                toggle_auto_exposure(int(input("Enter auto-exposure state (0: off, 1: on): ")))
            elif key == ord('f'):
                fov, amr = get_fov()
                print(f"Current FOV: {fov} mm, AMR: {amr}")
            elif key == ord('i'):
                set_flc_switch(int(input("Enter FLC Switch state (1: UV, 2: White, 3: Both, 4: Off): ")))
            elif key == ord('l'):
                set_flc_level(int(input("Enter FLC Level (1-32): ")))
            elif key == ord('b'):
                set_video_property("Brightness", int(input("Enter brightness value: ")))
            elif key == ord('r'):
                get_video_property("WhiteBalance")
            elif key == ord('c'):
                set_video_property("Contrast", int(input("Enter contrast value: ")))
            elif key == ord('w'):
                set_video_property("WhiteBalance", int(input("Enter WhiteBalance value: ")))
            elif key == ord('h'):
                set_video_property("Hue", int(input("Enter hue value: ")))
            elif key == ord('s'):
                set_video_property("Saturation", int(input("Enter saturation value: ")))
            elif key == ord('p'):
                set_video_property("Sharpness", int(input("Enter sharpness value: ")))
            elif key == ord('d'):
                apply_defaults()
            elif key == ord('g'):
                set_video_property("Gain", int(input("Enter gain value: ")))
            elif key == ord('q'):
                break

    listener.stop()
    camera.release()
    cv2.destroyAllWindows()

def main():
    microscope.SetVideoDeviceIndex(DEVICE_INDEX)
    microscope.Init()
    start_camera()
    apply_defaults()

if __name__ == "__main__":
    main()
```
